# Contributions and competition of Mg^2+^ and K^+^ in folding and stabilization of the Twister Ribozyme

**DOI:** 10.1101/2020.06.15.152744

**Authors:** Abhishek A. Kognole, Alexander D. MacKerell

## Abstract

Native folded and compact intermediate states of RNA typically involve tertiary structures in the presence of divalent ions such as Mg^2+^ in a background of monovalent ions. In a recent study we showed how the presence of Mg^2+^ impacts the transition from partially unfolded to folded states through a “push-pull” mechanism where the ion both favors and disfavors the sampling of specific phosphate-phosphate interactions. To better understand the ion atmosphere of RNA in folded and partially folded states results from atomistic Umbrella Sampling and oscillating chemical potential Grand Canonical Monte Carlo/Molecular Dynamics (GCMC/MD) simulations are used to obtain atomic-level details of the distributions of Mg^2+^ and K^+^ ions around Twister RNA. Results show the presence of 100 mM Mg^2+^ to lead to increased charge neutralization over that predicted by counterion condensation theory. Upon going from partially unfolded to folded states overall charge neutralization increases at all studied ion concentrations that, while associated with an increase in the number of direct ion-phosphate interactions, is fully accounted for by the monovalent K^+^ ions. Furthermore, K^+^ preferentially interacts with purine N7 atoms of helical regions in partially unfolded states thereby potentially stabilizing them. Thus, both secondary helical structures and formation of tertiary structures leads to increased counterion condensation, thereby stabilizing those structural features of Twister. Notably, it is shown that K^+^ can act as a surrogate for Mg^2+^ by participating in specific interactions with non-sequential phosphate pairs that occur in the folded state, explaining the ability of Twister to self-cleave at sub-millimolar Mg^2+^ concentrations.

## Introduction

RNA is currently a trending topic in fundamental biological studies and as a target for the development of therapeutic strategies (Disney 2019). Central to many functions and characteristics of RNA is its ability to fold into well-defined three-dimensional structures (Westhof and Auffinger 2000). Accordingly, it is important to understand the folding mechanisms of RNA. Over the years, various experimental and computational approaches have been used to investigate the folding patterns of RNAs (Onoa and Tinoco 2004; Šponer et al. 2018), including their interactions with ions (Heilman-Miller et al. 2001a; Heilman-Miller et al. 2001b; Kirmizialtin et al. 2012; Denesyuk and Thirumalai 2015; Panja et al. 2017). However, it is still unclear how the polyanionic structure of RNA in conjunction with the complex ionic atmosphere allows for RNA molecules to achieve stable tertiary structures.

It is well established that the negatively charged phosphate backbone of polynucleotides needs to overcome the huge electrostatic repulsion in order to fold into compact structures (Woodson 2010). The ionic atmosphere around the RNA provides the electrostatic shielding to facilitate this process (Draper 2004). The role of divalent ions to stabilize the folded tertiary structure of RNA molecules has been explored extensively (Li et al. 2011; Cunha and Bussi 2017; Šponer et al. 2018). Divalent ions, such as Mg^2+^, can be hundreds of times more efficient at facilitating and stabilizing the tertiary interactions as compared to monovalent ions like Na^+^ (Kim et al. 2002). However, a combination of mono- and divalent ions, such as found in physiological conditions, works in a synergistic way to further facilitate folding of RNA (Onoa and Tinoco 2004). For example, even with saturating Mg^2+^ there is a monovalent cation-binding site associated with the formation of the GACG RNA tertiary structure and Mg^2+^ is ineffective at this site (Shiman and Draper 2000). The Twister ribozyme represents an interesting case as it self-cleaves at Mg^2+^ at sub-millimolar concentrations (Panja et al. 2017; Korman et al. 2020), suggesting an important role of the monovalent ion in stabilizing folded states for this RNA.

Computational investigations of simultaneous mono- and divalent ion interactions with polynucleotides have observed limited success and have largely been performed on folded states or lack comprehensive spatial sampling of ions around the RNA (Yoo and Aksimentiev 2012; Ucisik et al. 2016; Fischer et al. 2018; Xi et al. 2018). A significant step forward are studies using coarse-grained models from the Thirumalai group that have had success in prediction of ion condensation, the specificity of Mg^2+^-RNA interactions and to understand RNA folding in mono- and divalent ion mixtures using both fully explicit ion representations and treatment of the monovalent ions using the reference interaction site model (RISM) (Denesyuk and Thirumalai 2013; Denesyuk and Thirumalai 2015; Denesyuk et al. 2018; Hori et al. 2019; Nguyen et al. 2019). A comparative study using small-angle x-ray scattering experiment and MD simulation was able to characterize the ionic cloud around a short RNA duplex (Pabit et al. 2010; Kirmizialtin et al. 2012). However, this approach used Rb^+^ as the monovalent ion and Sr^2+^ as the divalent ion diverting away from physiologically relevant ions. Overall, it is still not totally clear how an ionic atmosphere that includes both divalent and monovalent (Mg^2+^ and K^+^) ions recognizes and treats the conformational heterogeneity and flexibility along with polyanionic backbone of oligonucleotides and facilitates the folding of complex tertiary structures such as Twister ribozyme.

This study builds on our recent work where we have implemented a combination of enhanced sampling methods to understand the impact of Mg^2+^ ions at four different concentrations on partial unfolding of the Twister ribozyme (Kognole and MacKerell 2020). Umbrella sampling (US) was applied to model the energetics associated with sampling of the native and partially unfolded states of Twister ribozyme. The reaction coordinate (RC) was defined as the distance between centers of masses of two parts of the Twister ribozyme (Figure S1) such that with increasing distance the major tertiary interactions would be broken motivated by experimental studies on Twister (Panja et al. 2017). This yielded free energy profiles for different MgCl2 concentrations from the native states though partially unfolded states that lacked tertiary structure but maintained the secondary structural features; recent RNA folding studies have similarly shown secondary, helical structures to form prior to the formation of tertiary structures (Nguyen et al. 2019).

To investigate the impact of Mg^2+^ concentration on the free energy surfaces, Mg^2+^, K^+^ and Cl^-^ distributions were explicitly modeled along the RC using oscillating μ_ex_ Grand Canonical Monte Carlo / Molecular Dynamics (GCMC/MD) (Lakkaraju et al. 2014). The GCMC/MD approach was previously shown to recapitulate known Mg^2+^ binding sites as well as identify new sites in four RNAs (Lemkul et al. 2016) and the approach facilitated a study of Mg^2+^ binding to and allosteric modulation of the μ-opioid receptor (Hu et al. 2019; MacKerell 2019). Overall, the combined US-GCMC/MD approach achieved resampling of both mono- and divalent ions around the native and partially unfolded states of Twister yielding an atomic level picture of how Mg^2+^ contributes to the folding of Twister. Specifically, the study showed the divalent ion to stabilize RNA through overall stabilization of specific non-sequential non-bridging phosphate oxygen (NBPO) pairs, consistent with the observation of Hori et al. (Hori et al. 2019). Notably, this was observed to occur through a “push-pull” mechanism where specific non-sequential NBPO pairs were stabilized by Mg^2+^ while other NBPO pairs were simultaneously destabilized. Presently, further analyses of those simulations are presented revealing atomic details of the ionic atmosphere around Twister from native to partially unfolded states of the RNA, including the nature of the competition between the divalent Mg^2+^ and monovalent K^+^ counterions included in the study.

## Results and Discussion

The previous study focused on local interactions of Mg^2+^ with the RNA and how they stabilize the native state and impact free energy of the sampled partially unfolded states. From that study a “push-pull” mechanism was established in which the presence of Mg^2+^ leads to increased sampling of some short interactions between NBPOs while decreases in the sampling of other short NBPOs simultaneously occur. This leads to an overall increase in the sampling of short non-sequential NBPO interactions, thereby favoring the folded state. As those simulations were performed in a background of 100-200 mM KCl at four different Mg^2+^ concentrations (Table S1) significant atomistic information on the ion atmosphere around the RNA along the sampled folding profile is available. Here, we present a detailed analysis of the ion distribution around the Twister ribozyme at the fully folded, inflection point and the partially folded structures sampled in the US simulations in the previous study. The potentials of mean force from the US calculations are shown in Figure S1b, which indicates regions of the three regions along RC that are referred in the following analyses.

### Impact of Mg^2+^ on charge neutralization

CIC theory has been widely used towards characterization of the ion atmosphere around nucleic acids (Heilman-Miller et al. 2001b), although it is formally only applicable to polynucleotides such as DNA duplexes that have a rod-like shape (Savelyev and MacKerell 2014). The primary role of the counterions is neutralization of the polyanionic charge allowing the oligonucleotide to collapse into compact structures (Thirumalai et al. 2001; Draper et al. 2005). Accordingly, initial analysis investigated the effect of different concentrations of divalent ions around the RNA on percent charge neutralization (%CN) (Savelyev and MacKerell 2014). According to CIC theory for a rod-like structure with uniformly distributed charge there will be 76% charge neutralization by condensed counterions within a 9 Å Manning radius (Manning 1978). Although Twister has a more complex structure than the rod-like shape of duplex DNA, we calculated the average number of counterions condensed around the non-hydrogen atoms of Twister to find the distance needed to achieve a target charge neutralization of 76% and how the increasing Mg^2+^ concentration impacted the charge neutralization. Figure 1 shows that at the native state with increasing Mg^2+^ concentrations the distance to achieve 76% charge neutralization reduced significantly from ~ 11 Å at 0 mM MgCl_2_ to ~ 5.5 Å at 100 mM MgCl_2_. For the 10 mM and 20 mM MgCl_2_ systems the relevant distances are in the range of 9 to 10 Å. The 5.5 Å distance for 76% charge neutralization in the 100 mM system indicates the effectiveness of Mg^2+^ ions to condense around the RNA. This is obviously associated with the divalent nature of Mg^2+^ ion in conjunction with its high charge density requiring a lower entropic costs for redistribution of ions and waters around the complex RNA (Draper 2004). This indicates that Mg^2+^ at the higher concentration contributes to stabilization of the RNA via charge neutralization in addition to specific interactions with the RNA seen in our previous work (Kognole and MacKerell 2020) and earlier by Hori et al (Hori et al. 2019).

**Figure 1.**
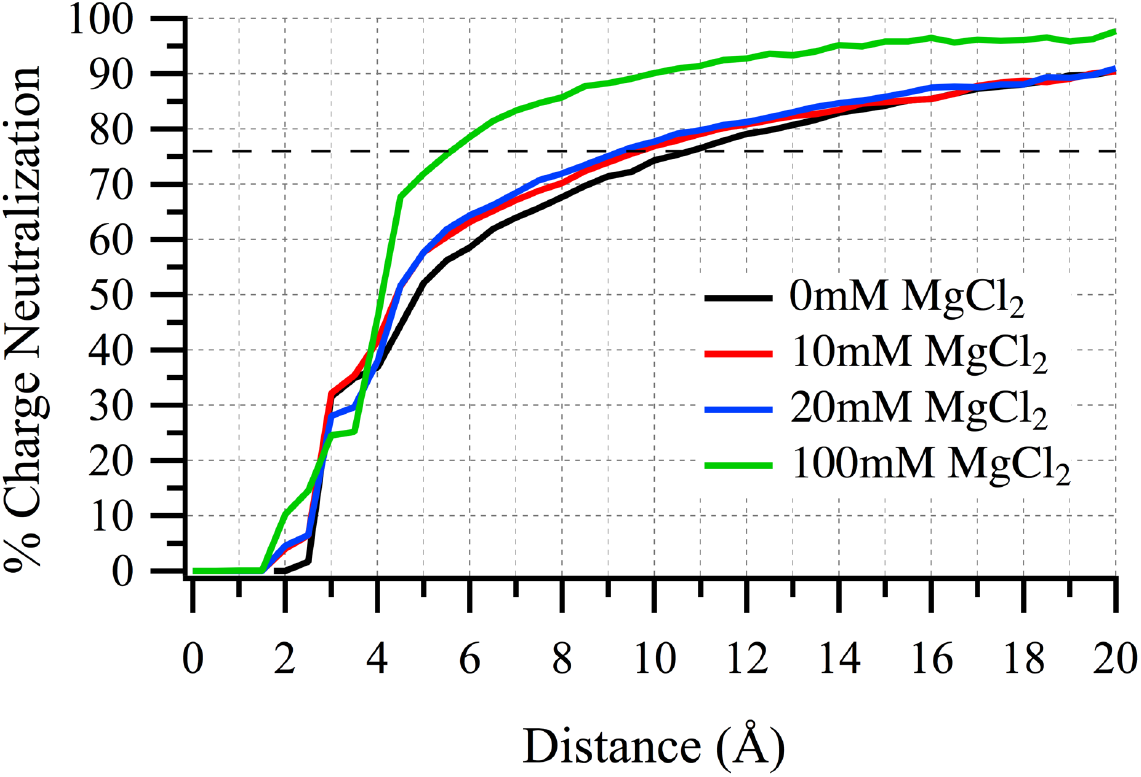
Percent charge neutralization by counterions at the native state (RC = 14 – 16 Å) at different MgCl_2_ concentrations as a function of distance from all non-hydrogen atoms of Twister. The dashed line indicated the 76% charge neutralization predicted by counterion condensation theory (see text).

### Increased ion condensation near the folded state

The ion distributions and RNA conformations from the US allows investigation of the effect of counterion condensation on charge neutralization at various partially folded states of Twister ribozyme. Figure 2 shows the %CN based on an 8 Å cutoff as a function of the RC. In all systems there is approximately a 4 – 5% reduction in charge neutralization at the most unfolded conformations sampled compared to the native state. As reported in our previous study, even in the most unfolded states at RC = 40 Å the amount of secondary structure is similar to that in the native state. Thus, the presence of decreased charge neutralization at longer RC values indicates the contribution of the tertiary structure of RNA on the ability of the counterions to condense around the RNA, which is not anticipated in CIC theory, as previously discussed (Hori et al. 2019). This effect occurs with K^+^ as well as Mg^2+^ as indicated by its occurrence in 0 mM MgCl_2_. The results also indicate that the condensation of K^+^ contributes to stabilization of the folded states of Twister, consistent with the ability of the RNA to assume compact structures in low Mg^2+^ concentrations.

**Figure 2.**
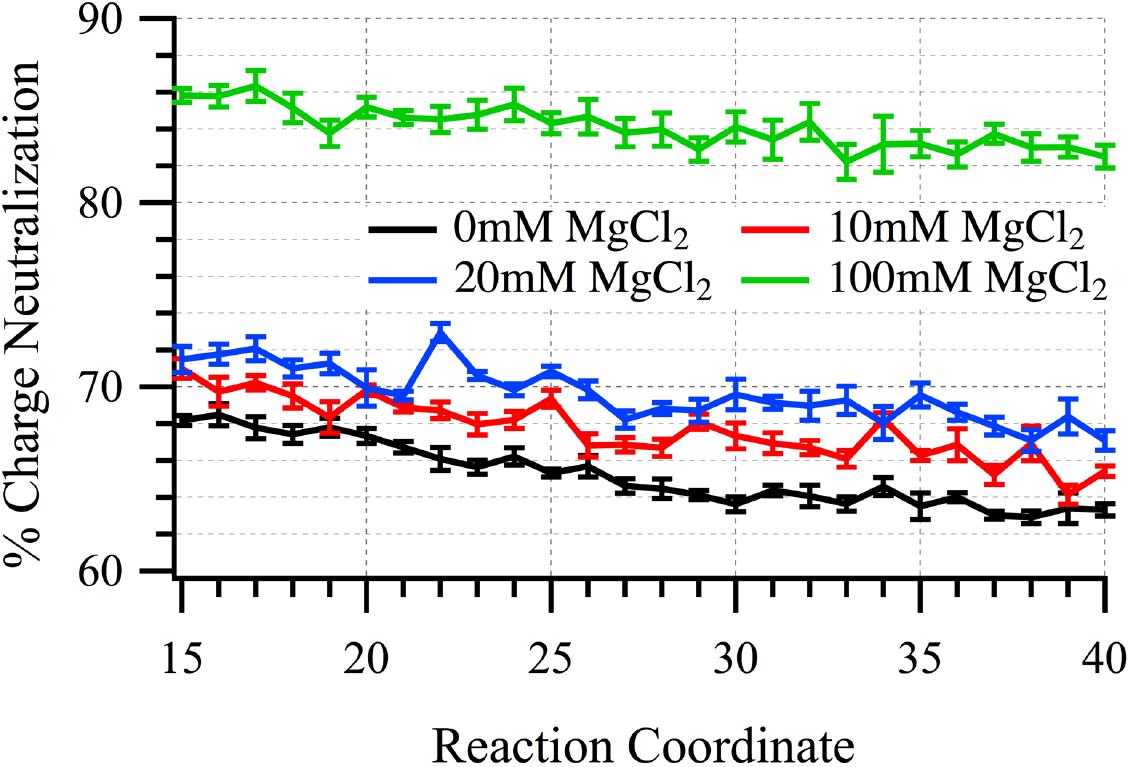
Percent charge neutralization (%CN) based on an 8 Å cutoff from all heavy atoms of twister at various stages of the reaction coordinate.

Given the increased %CN as the RNA assumes the folded state more detailed analysis was undertaken by determining the number of ions in the environment of the RNA along the sampled RC. The analysis involved different cutoffs presented in the methods where a 3 Å cutoff defines direct contact of ions with NBPOs, 3 – 5.5 Å represents outer-shell contacts, 5.5 – 8 Å range represents diffuse (non-dehydrated) contacts and ions beyond 8 Å are not considered as condensed ions. As shown in Table 1, for the 100mM MgCl_2_ system the number of both Mg^2+^ and K^+^ in direct contact with the RNA increase upon going from partially unfolded to more folded states. Interestingly, with Mg^2+^ the increase of ions directly interacting with the NBPOs is associated with a decrease in the number of outer-shell ions with the number of non-dehydrated ions (Mg^2+^ within 5.5 - 8 Å) relatively low and approximately constant across the reaction coordinate. Notably, the overall number of Mg^2+^ in and outside of the condensed layer, as defined by an 8 Å cutoff, is constant along the RC. In contrast, with K^+^ the increase in direct RNA-ion interactions is associated with an increase in the number of outer-shell ions leading to an overall increase in the number of K^+^ in the condensed layer while the number of non-condensed ions decreases. Table S2 includes number of K^+^ ions in the 0, 10 and 20 mM MgCl_2_ systems which may be compared to the 100 mM MgCl_2_ results in Table 1. As expected, the decreased or absence of Mg^2+^ in those systems leads to a significant increase in the number of K^+^ ions near that RNA as required to neutralize the polyanionic RNA with the total increased number of ions due to the higher concentration of K^+^ in those systems (Table S1). Notably, the trend where the number of both direct and outer shell K^+^ ions increases associated with an overall increase in the number of ions in the condensed layer occurs similar to that at 100 mM MgCl_2_, with the magnitude in the change being larger (~4 for 0 MgCl_2_ vs. ~2). Thus, the folding of the RNA leads to more direct ion-RNA interactions with both Mg^2+^ and K^+^ though the pool for those additional ions differs, with the increased charge neutralization being associated with increased condensation of the monovalent K^+^ in both the presence and absence of divalent Mg^2+^.

**Table 1.** Number of ions around the phosphate backbone of Twister in 100mM MgCl_2_. The 3 Å cutoff corresponds to direct contact of ions with non-bridging phosphate oxygens (NBPOs), 3 – 5.5 Å range represents outer-shell contacts, 5.5 – 8 Å range represents diffuse (non-dehydrated) contacts. The 8 Å cutoff defines the full condensed-ion atmosphere. The errors represent standard error of mean calculated over five 12 ns trajectories.

| **Mg<sup>2+</sup> within** | **0 – 3 Å** | **3 – 5.5 Å** | **5.5 – 8 Å** | **0 – 8 Å** | **beyond 8 Å** |
| --- | --- | --- | --- | --- | --- |
| **RC_15-18** | 2.30 ± 0.27 | 13.91 ± 0.34 | 1.66 ± 0.29 | 17.87 ± 0.21 | 22.13 ± 0.21 |
| **RC_19-22** | 2.50 ± 0.35 | 13.58 ± 0.42 | 1.71 ± 0.33 | 17.79 ± 0.23 | 22.21 ± 0.23 |
| **RC_23-26** | 2.36 ± 0.25 | 14.42 ± 0.45 | 1.71 ± 0.52 | 18.49 ± 0.36 | 21.51 ± 0.36 |
| **RC_27-30** | 1.86 ± 0.40 | 14.19 ± 0.56 | 1.71 ± 0.55 | 17.76 ± 0.39 | 22.24 ± 0.39 |
| **RC_31-34** | 1.71 ± 0.38 | 14.75 ± 0.53 | 1.61 ± 0.47 | 18.07 ± 0.29 | 21.93 ± 0.29 |
| **RC_35-40** | 1.69 ± 0.27 | 14.73 ± 0.40 | 1.70 ± 0.40 | 18.12 ± 0.27 | 21.88 ± 0.27 |
| **K<sup>+</sup> within** | **0 – 3 Å** | **3 – 5.5 Å** | **5.5 – 8 Å** | **0 – 8 Å** | **beyond 8 Å** |
| **RC_15-18** | 1.91 ± 0.14 | 6.49 ± 0.32 | 1.08 ± 0.45 | 9.48 ± 0.34 | 32.52 ± 0.34 |
| **RC_19-22** | 1.53 ± 0.13 | 6.78 ± 0.24 | 1.02 ± 0.30 | 9.33 ± 0.22 | 32.67 ± 0.22 |
| **RC_23-26** | 1.01 ± 0.12 | 5.66 ± 0.39 | 1.06 ± 0.51 | 7.73 ± 0.35 | 34.27 ± 0.35 |
| **RC_27-30** | 1.07 ± 0.09 | 6.04 ± 0.40 | 0.98 ± 0.59 | 8.09 ± 0.44 | 33.91 ± 0.44 |
| **RC_31-34** | 0.94 ± 0.07 | 6.03 ± 0.28 | 1.12 ± 0.40 | 8.09 ± 0.29 | 33.91 ± 0.29 |
| **RC_35-40** | 0.97 ± 0.11 | 5.65 ± 0.30 | 1.08 ± 0.38 | 7.70 ± 0.26 | 34.30 ± 0.26 |

Recently, it was reported that one inner-shell coordination with divalent ion releases two monovalent ions from the ionic atmosphere, a process that is entropically favorable (Nguyen et al. 2019). In the present study, based on ion counting data, upon going from the 0 mM to 100 mM MgCl_2_ systems the average number of K^+^ ions released from the local ion atmosphere based on ions in direct contact with the RNA is close to 2.0 (Table 2, 0 – 3 Å range) consistent with the RISM study. However, if the full condensed layer of ions is considered (0 – 8 Å range) then the number released is smaller, being approximately 1.3 to 1.4. In the 10, 20 and 100 mM MgCl_2_ systems these values are 1.65, 1.56 and 1.32 K^+^ ions, respectively, for the full condensed layer of ions based on the 0 – 8 Å range (Supplementary Information Table S3). These results show that while each Mg^2+^ in direct contact with the RNA displaces two K^+^ ions, some monovalent ions still are in direct contact with the RNA through interactions with the nucleobases as discussed below. Overall, the present results indicate that Mg^2+^ can displace K^+^ from direct interactions at a ratio of 1:2 as previously shown, but the extent of displacement from the overall condensed phase is lower indicating that K^+^ contributes to the stabilization of the RNA in the presence of Mg^2+^.

**Table 2.** Average number of K^+^ displaced per Mg^2+^ ions for various cutoffs at different stages of reaction coordinate for 100 mM MgCl_2_ system compared to 0 mM MgCl_2_ system.

|  | 0 – 3 Å | 3 – 5.5 Å | 0 – 5.5 Å | 5.5 – 8 Å | 0 – 8 Å |
| --- | --- | --- | --- | --- | --- |
| **RC_15-18** | 2.03 | 1.24 | 1.35 | 1.78 | 1.39 |
| **RC_19-22** | 1.78 | 1.23 | 1.32 | 1.75 | 1.36 |
| **RC_23-26** | 1.72 | 1.22 | 1.29 | 1.58 | 1.32 |
| **RC_27-30** | 2.02 | 1.19 | 1.28 | 1.62 | 1.32 |
| **RC_31-34** | 2.53 | 1.10 | 1.25 | 1.63 | 1.28 |
| **RC_35-40** | 2.30 | 1.14 | 1.26 | 1.50 | 1.28 |

### Competition between Mg^2+^ and K^+^ for non-sequential NBPO pairs

A collection of six non-sequential NBPO pairs need to assume close interactions to allow Twister to assume the folded state (Figure S2). Visualization of the ion distributions around the full folded state of Twister in the form of GFE maps is shown in Figure 3 at the four Mg^2+^ concentrations. At 100 mM MgCl_2_ the Mg^2+^ dominates the local environment around the RNA including the vicinity of the important non-sequential NBPO, as detailed in our previous study (Kognole and MacKerell 2020). However, consistent with the data in Table 1, K^+^ occupies the region adjacent to the RNA even though there is an additional pool of Mg^2+^ beyond the condensed layer around the RNA available to compete with the K^+^. In the absence of Mg^2+^ ions, the K^+^ ions occupy the space around both the negatively charged backbone as well as the nucleobases. The distributions show that with decreasing Mg^2+^ those ions sample local regions adjacent to the specific non-sequential NBPO pairs. These results show that the local ion atmosphere of the RNA is comprised of a combination of mono- and divalent ions even in the presence of high concentrations of Mg^2+^. Thus, while coordination of the Mg^2+^ ion at specific binding sites facilitates stabilization of the RNA, the monovalent ions are not totally excluded from the regions adjacent to the RNA.

**Figure 3.**
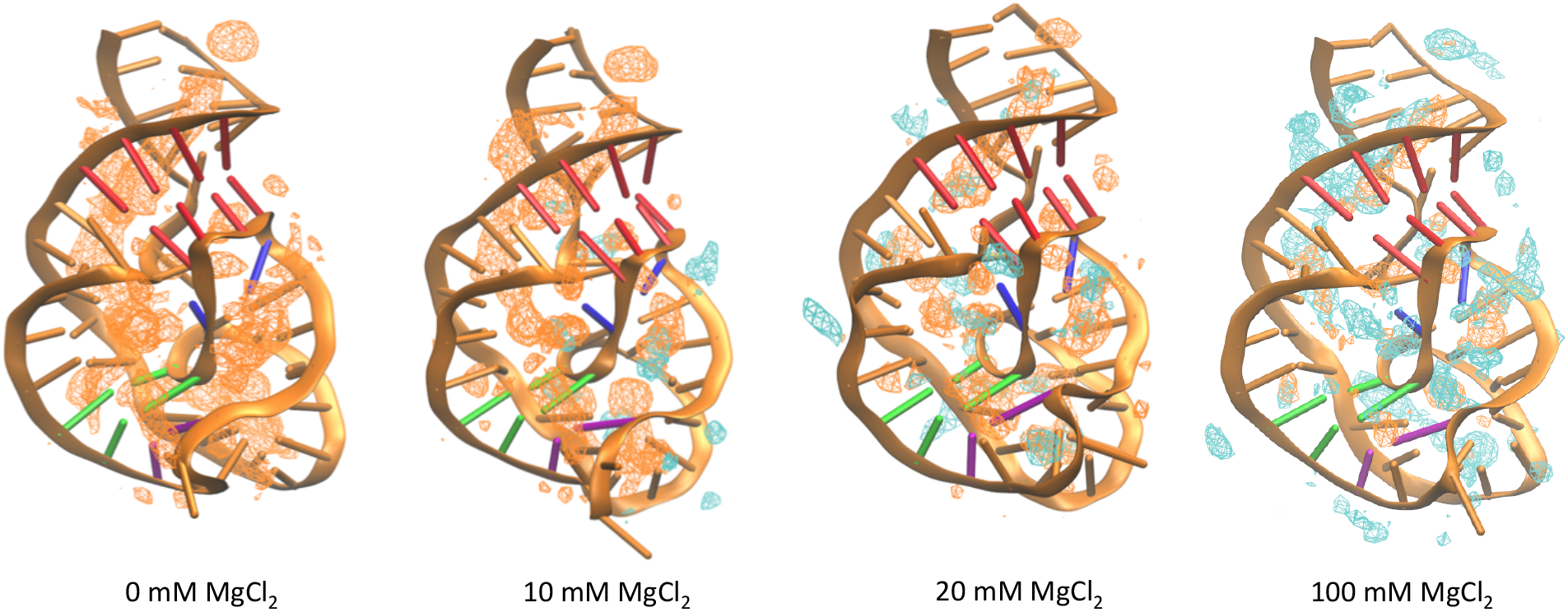
GFE maps for Mg^2+^ (cyan mesh) and K^+^ (orange mesh) distributions around the native state of Twister at different Mg^2+^ concentrations. The GFE maps are contoured at −2.5 kcal/mol.

Our previous study of non-sequential NBPO pairs and their interaction with Mg^2+^ ions showed that whenever the NBPO pair has a distance lower than 9 Å there is significant probability of Mg^2+^ being in contact with both the phosphates (Kognole and MacKerell 2020). Such interactions contribute to the formation of tertiary interactions leading to assumption of the native state. The consistently higher contacts with Mg^2+^ at certain close NBPO pairs, such as region around NBPO pair 4-46 in the P1 helix (Figure S2), contributed to the push-pull stabilization of the ribozyme that facilitate formation of the T1 and other tertiary interactions. To investigate how effectively K^+^ can act as a surrogate for Mg^2+^ analysis of the probability of finding K^+^ ions around these close non-sequential NBPO pairs present in the tertiary structure was performed along the RC.

Presented in Figure 4 are the probabilities of forming the short NBPO pairs and K^+^ being within 6.5 Å of NBPOs of both phosphates along the RC at both 0 and 100 mM MgCl_2_. In high Mg^2+^ the probability of K^+^ being close to both NBPOs is 0.2 or less with the exception of the C20-G30 pair between RC = 24 and 31 Å (red dashed lines). However, in all cases in the absence of Mg^2+^ the probability of K^+^ being close to both NBPOs is significantly higher (black dashed lines). Notably, along the entire RC higher probabilities of K^+^ interacting with both NBPOs correlate with the NBPOs being close, as required to assume the folded state. The overall pattern of short NBPO pairs are similar in both 0 and 100 mM MgCl_2_ with the probabilities in the presence of Mg^2+^ typically being higher consistent with its stabilizing effect (red solid lines). However, exceptions exist, including PP pair 9-28 from the inflection point to the folded state and with PP pair 32-41 in the region of 20 – 25 Å of the RC, consistent with the push-pull mechanism. These results indicate the ability of K^+^ to act as an effective surrogate for Mg^2+^ with Twister through simultaneous interactions of both phosphates in the case of short NBPO pairs. Based on these results it is evident that K^+^ can stabilize phosphate-phosphate interaction through site-specific interactions though it is not as effective as Mg^2+^.

**Figure 4.**
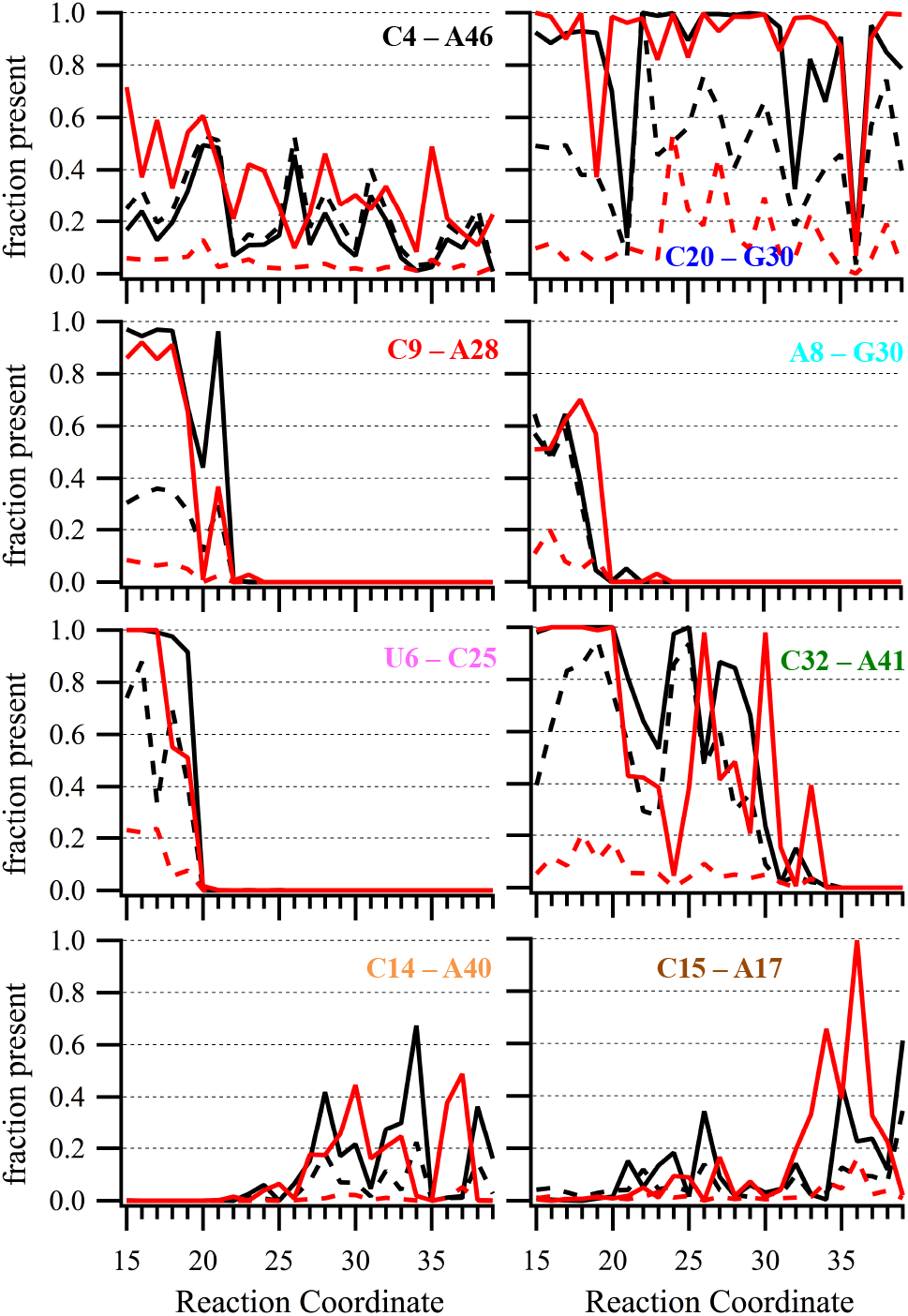
Adjacent non-bridging phosphate oxygen (NBPO) pairs in contact with K^+^ ions at 0 mM MgCl_2_ (black lines) and at 100 mM MgCl_2_ (red lines). Shown is the probability of selected NBPO pairs within 9 Å (solid lines) and the probability of K^+^ ions (dashed lines) within 6.5 Å of NBPOs of both phosphates.

Experimentally, it has been shown that sub millimolar concentrations of Mg^2+^ ions are sufficient to nucleate the folding of ribozymes in the presence of buffer containing K^+^ (Denesyuk and Thirumalai 2015), and even lower concentrations to self-cleave in case of Twister (Panja et al. 2017; Korman et al. 2020). As Twister is largely folded at 20 mM Mg^2+^, where there is a total of 8 ions in the simulation system, the probability of K^+^ vs Mg^2+^ interacting with the NBPO pairs was compared at 20 and 100 mM MgCl_2_ (Figure 5). As expected at 100 mM MgCl_2_, the probability of K^+^ being adjacent to the NBPO pairs is generally significantly lower than for the divalent Mg^2+^ ion, though K^+^ is present in all cases (Figure 5b). In addition, there are cases where the probability of K^+^ is similar or higher than that of Mg^2+^ including with the PP pair 20-30 after the inflection point (RC = 20) and 6 - 25 NBPO pair where the probability of Mg^2+^ versus K^+^ sampling is similar near the native state. However, at 20 mM MgCl_2_ (Figure 5a) K^+^ sampling adjacent to the nonsequential NBPOs dominates over Mg^2+^. This is evident for the majority of the short NBPO pairs, including near the native state and along the majority of the RC, showing that K^+^ ions occupy the same binding sites as Mg^2+^ in a complementary fashion. Based on these results it appears that in the presence of a limiting amount of Mg^2+^ ions, K^+^ ions can play a crucial supporting role by facilitating the sampling of short NBPO pairs while available Mg^2+^ ions may have already occupied other sites. This phenomenon is suggested to allow for the experimentally observed folding of Twister at low Mg^2+^ concentrations as well as to self-cleave at even lower concentrations.

**Figure 5.**
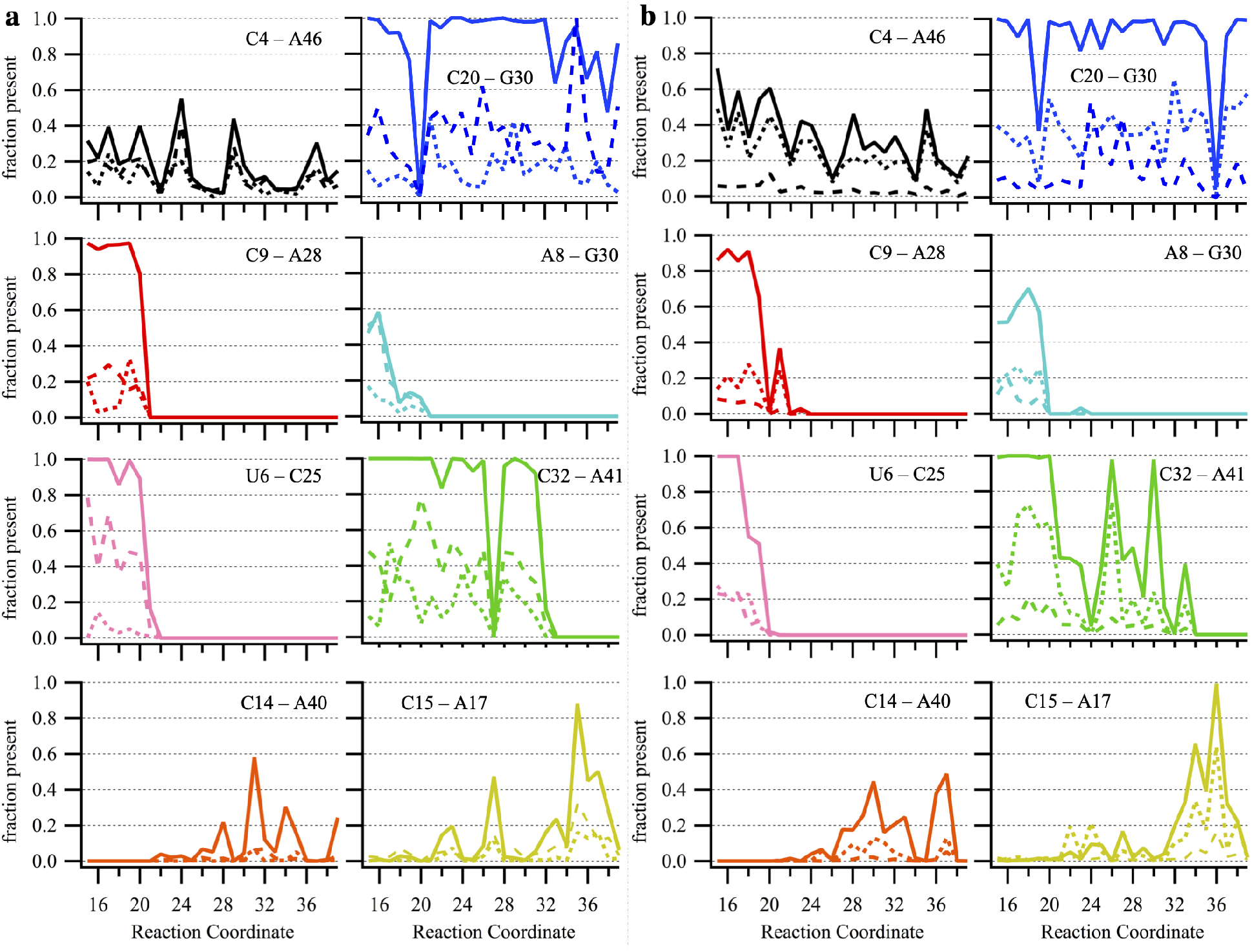
Adjacent NBPO pairs in contact with Mg^2+^ and K^+^ ions at a) 20mM and b) 100 mM MgCl_2_. Shown is the probability of selected NBPO pairs within 9 Å (solid line) and the probability of Mg^2+^ (dotted line) or K^+^ ions (dashed line) within 6.5 Å of NBPOs of both phosphates.

### Ion-RNA backbone and base interactions as a function of folding

Beyond interactions with the native fold of RNA, ions can potentially interact with the nucleobases of the RNA as well as with the phosphates in partially unfolded states, thereby facilitating the folding process. Previously, a coarse-grained study has reported a relationship between K^+^ ion condensation and residual helical structures in unfolded ribozyme at low Mg^2+^ concentrations (Hori et al. 2019). The GCMC ion sampling method in conjunction with the use of US to sample partially unfolded states allows for a comprehensive atomistic picture of the ion atmosphere to be obtained as a function of the extent of folding. A general picture of ion condensation associated with RNA folding is shown in Figure 6 showing the changes in Mg^2+^ and K^+^ ion distribution around the Twister at various stages of the RC in the 20mM MgCl_2_ system. Similar analysis at the 0, 10 and 100 mM concentrations is provided in SI Figures S3, S4 and S5. The distributions in Figure 6 show that K^+^ is broadly distributed around the RNA including in the vicinity of the nucleobases of helical regions. Additionally, K^+^ is close to the phosphodiester backbone and in the buried region between the Twister catalytic site and T2 tertiary interaction (Figure S2). At RC = 21 & 24, around the inflection point and just after it, the distribution of K^+^ around the phosphodiester backbone is dissipating. And with further unfolding K^+^ is predominantly interacting with nucleobases of helical regions.

**Figure 6.**
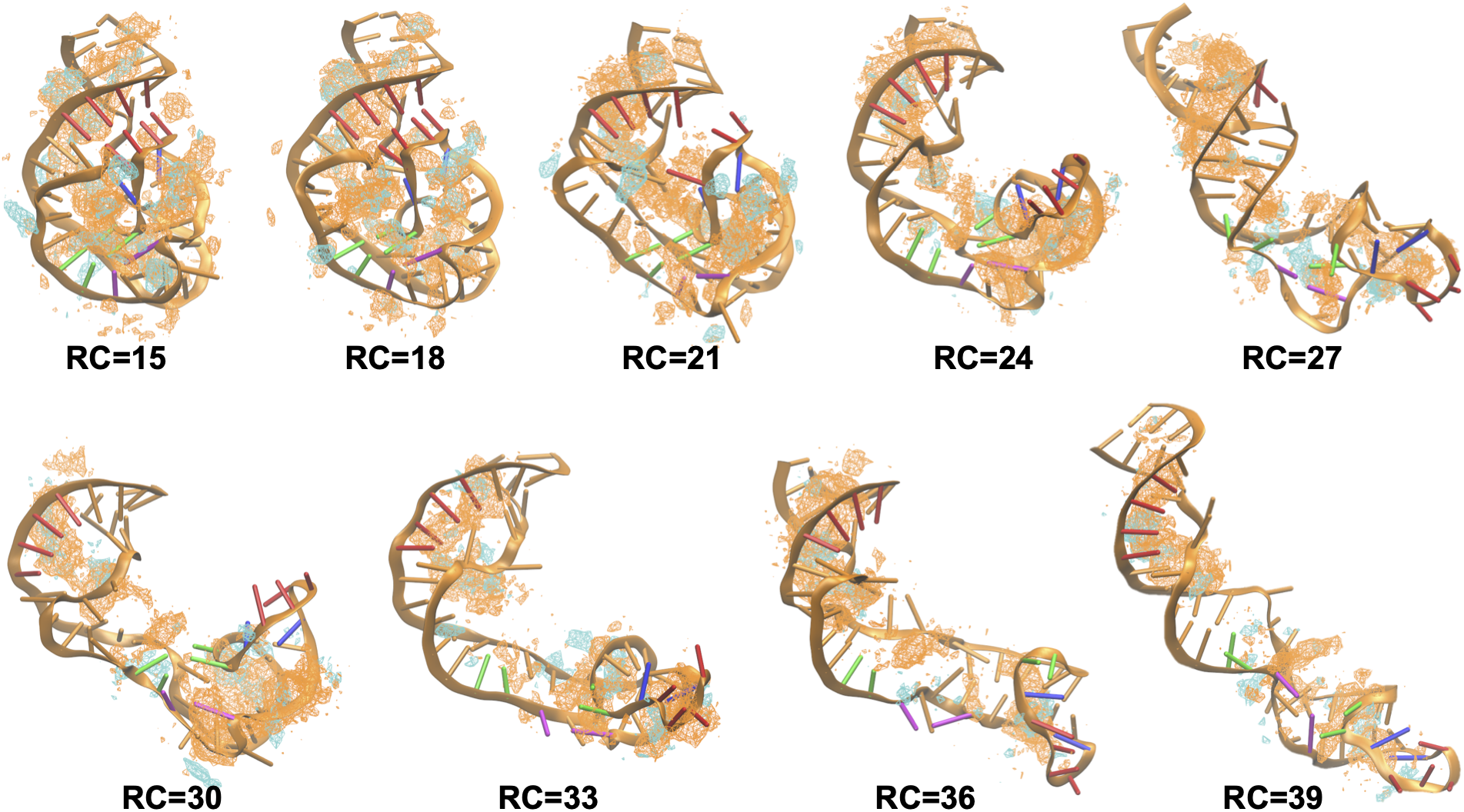
GFE maps of the distribution of K^+^ (orange) and Mg^2+^ (cyan) ions around the Twister ribozyme at various stages along the umbrella sampling reaction coordinate in 20 mM MgCl_2_. Twister is shown as orange cartoon with T1 bases in red and T2 bases in green cartoon. GFE level is −2.0 kcal/mol.

Further analysis focused on understanding how the two types of ions compete for the different classes of functional groups comprising the RNA along the RC. Analysis involved the RDFs of ions around the NBPOs, the Gua-N7/Ade-N7 atoms and the O2/O4/O6 atoms of the nucleobases at three regions along RC; the native state, the inflection region and the partially unfolded state at 39 Å (Figure S1b). As previously shown, Mg^2+^ dominates the direct interactions with the NBPOs with only minimal direct interactions with the nucleobases (Figure S6) (Cunha and Bussi 2017; Šponer et al. 2018). The lack of direct interactions of Mg^2+^ ions with N7 has been previously reported (Leonarski et al. 2016) which has been suggested to be due to N7 not providing enough electrostatic attraction for dehydration of inner-shell water molecules (Peschke et al. 1998). Similar contributions likely lead to the lack of direct Mg^2+^ interactions with the oxygens atoms of the nucleobases consistent with a PDB survey of DNA showing that nucleobase carbonyl groups are poor binders with respect to direct Mg^2+^ inner-shell interactions but participate in direct interactions with monovalent ions (Leonarski et al. 2019). This pattern is maintained at the three regions of the RC shown in Figure S6, with direct interactions dominating with the phosphate oxygens and the outer-layer interactions dominating with the nucleobases. Thus, the impact of Mg^2+^ on RNA stabilization is not associated with interactions with the nucleobases to a significant degree and the pattern of interactions is not significantly altered by the extent of folding of the RNA.

In the case of K^+^ more interesting behavior in the RDFs is observed (Figure 7). With the NBPOs as well as the nucleobases the number of direct interactions are significant. Of these the interactions with N7 of the purines are the most prominent. In particular, Gua N7 has multiple factors contributing to direct interactions with monovalent ions. These include accessibility to ions in helical structures, especially in the major groove of DNA, favorable electrostatic potential and a supporting role of O6 that makes it a more effective binding site (Lippert 2000). The magnitude of the direct inner-layer interactions at the phosphate NBPOs and the nucleobase oxygens is similar with more outer-layer interactions occurring with the anionic phosphate backbone. More significant is the trend showing the number of direct interactions with the phosphate NBPOs and N7 atoms to systematically decrease upon going from the folded to the partially unfolded state. This is supported by the ion counting data shown in Table S2 for the NBPOs and is associated with more of the monovalent ion moving into the condensed layer upon folding, as discussed above. This pattern indicates that the direct interactions of K^+^ with the N7 of Gua and Ade as well as with the phosphate oxygens contribute to stabilizing the folded state of Twister.

**Figure 7.**
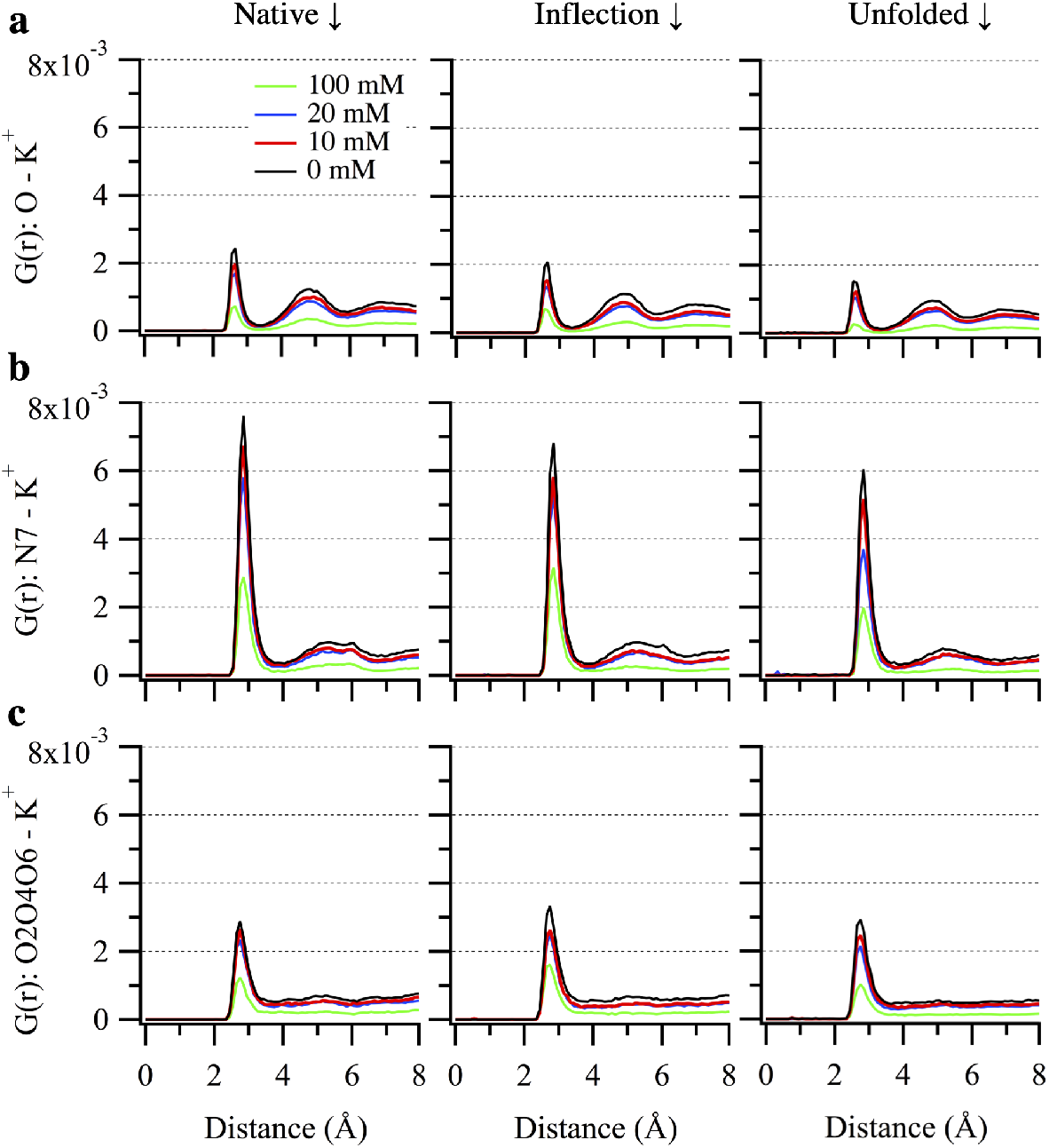
Radial distribution functions of K^+^ around a) the non-bridging phosphate oxygens (NBPO), b) the Gua-N7/Ade-N7 atoms, and c) O2/O4/O6 atoms of the nucleobases at native state (RC=14-16), inflection point (RC=19-21) and unfolded state (RC=38-40).

### Ion interactions with helical versus non-helical regions

To evaluate the impact of helical structure on the interactions of ions with the Twister RNA, we undertook analysis of ion-RNA interactions in helical and non-helical regions. The helical region corresponds to 26 nucleotides that participate in WC base-pairs of P1, P2 and P4 helices in the native state (Figure S2a). The remainder of the 28 nucleotides comprise the non-helical regions. RDFs of Mg^2+^ and K^+^ were calculated around NBPOs and the coordinating atoms (N7 and O2/O4/O6) of nucleobases at 100 mM MgCl_2_ at the three stages along RC (Figure 8). In the case of interactions with the NBPOs, both Mg^2+^ and K^+^ participate in a greater level of direct inner-shell coordination in the non-helical regions. The trend is more pronounced with K^+^ with a substantial decrease in the direct interactions occurring in the unfolded states. A similar but smaller decrease occurs with Mg^2+^. With the nucleobases substantial differences occur between Mg^2+^ and K^+^. With both N7 and the nucleobase oxygens direct interactions occur with Mg^2+^ in the non-helical regions, though the levels are much lower than with K^+^. In addition to the greater number of direct nucleobase-K^+^ interactions there are significantly more such interactions with the helical states. In combination with the results above, with K^+^ the increased direct interactions in the helical regions indicate that the monovalent ion contributes to stabilization of the helical regions. This is consistent with a report indicating that the stability of G-C base pairs is enhanced by binding of monovalent ions at the N7 position through polarization (Šponer et al. 2000) as well as studies showing that monovalent ions stabilize the helical, secondary interactions of nucleic acids (Shiman and Draper 2000; Draper 2008), while the results contrast remarks from Leonarski et. al. (Leonarski et al. 2016) indicating purine N7 binding sites are “not of primary importance in RNA and DNA.” In the case of Mg^2+^ the small, but increased level of direct interactions in the nonhelical states suggests that the lack of secondary structure leads to increased accessibility to the bases allowing for a small number of interactions that are not accessible in compact folded states.

**Figure 8.**
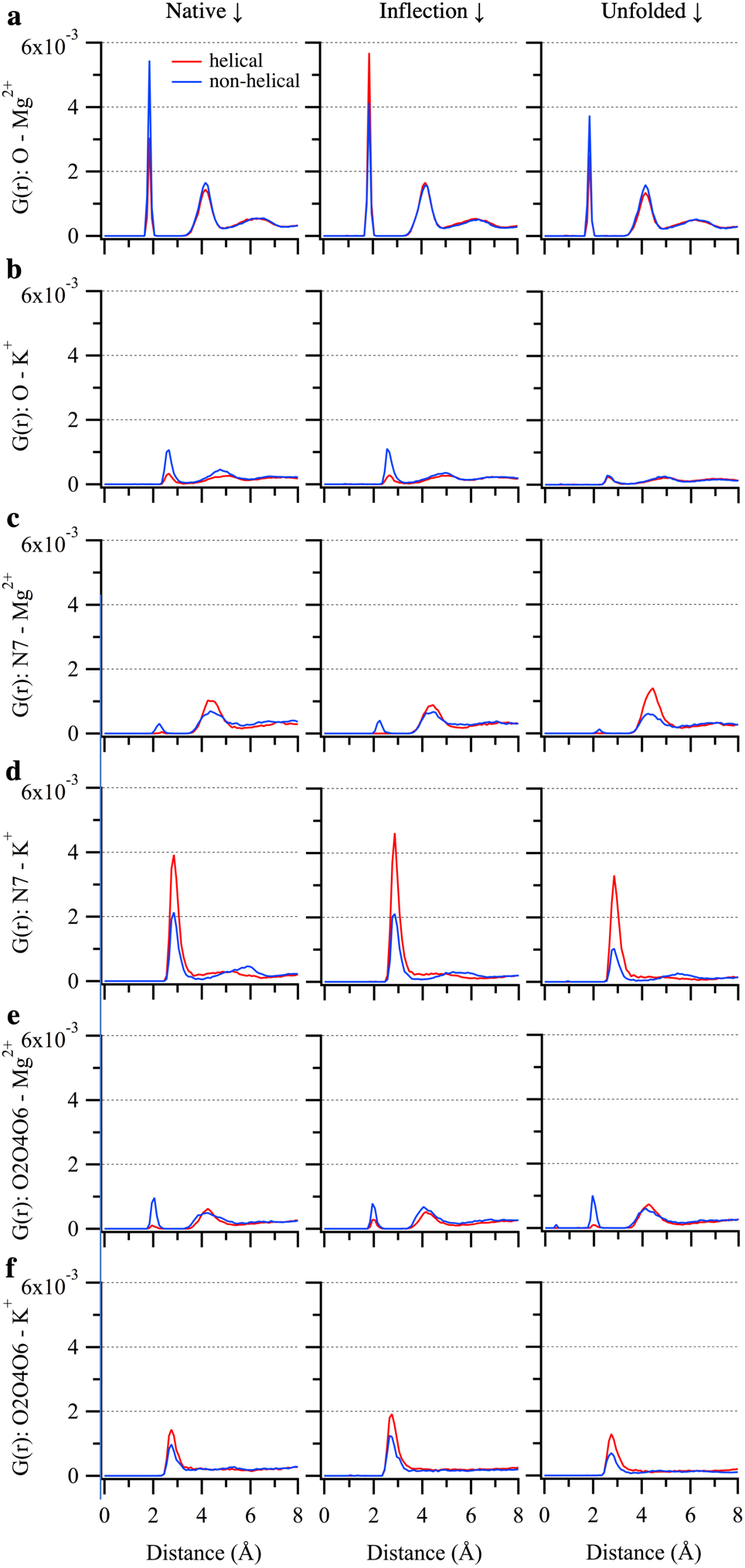
Radial Distribution Functions (RDF) of Mg^2+^ (a, c, e) and K^+^ (b, d, f) around NBPOs, Ade-N7/Gua-N7, and O2/O4/O6 atoms of nucleobases that participate or don’t participate in helical structures, respectively. From left to right the plots correspond to RDFs calculated at native state (RC=14-16), inflection point (RC=19-21) and unfolded state (RC=38-40) for 100mM MgCl_2_. Red line – Helical region, Blue line – Non-helical region.

Concerning differences between the folded, inflection and partially unfolded state some changes are observed. With Mg^2+^ differences between the three states are generally minimal with the largest difference being an increase in the outer-layer coordination of N7 in the helical regions of the unfolded state. This further suggests that the overall interactions between RNA and Mg^2+^ are facilitated by the lack of secondary and tertiary structure, with the specific interactions related to close non-sequential NPBO pairs being an exception. With K^+^ the trend towards decreased interactions in the more unfolded state is evident for the NBPOs and the nucleobases. This occurs with interactions with the nucleobases in both the helical and non-helical regions while the presence of direct interactions with the phosphates occurring in the native and inflection states with the non-helical regions are lost in the unfolded states. These changes and the increased sampling of direct inner-shell interactions in the helical state further indicate the contribution of the monovalent ion to stabilization of the helices with the contribution increasing as the RNA assumes more folded conformations.

An interesting observation with K^+^ is the increase in the direct interactions with the N7 and O2/O4/O6 atoms of the nucleobases. To investigate this more closely ion counting based on a 3.5 Å cutoff was undertaken over all the nucleobase coordinating atoms. For all four systems there is a decrease in the count in the unfolded state in both helical and non-helical regions. However, in the case of non-helical regions the total decrease in count from native to unfolded is larger (~1.0) than in the helical regions (~0.5). With the decreasing Mg^2+^ concentration, higher number of K^+^ ions are available to interact with the nucleobases in both helical and non-helical regions. The average total increase from 100 mM to 0 mM system is bigger for non-helical region (~4.0) as compared to helical region (~2.5). This overall increase in direct interactions of K^+^ ions with nucleobases in the absence of Mg^2+^ is consistent with RDFs in Figure 7. These differences further indicate the role of K^+^ in stabilizing Twister through direct interactions with the RNA by compensating for the lack of Mg^2+^.

In conclusion, the present study takes advantage of a unique combination of umbrella sampling with GCMC/MD ion sampling in the context of an all-atom model allowing detailed, atomic level analysis of the ionic atmosphere and specific ion-RNA interactions of the Twister ribozyme for both folded and partially unfolded states. This capability offers an additional approach along with the advances in coarse grain models to study ion-RNA interactions. Results show that monovalent ion binding sites in RNA are highly dependent on concentration of divalent ions. The RDF analysis of mono- and divalent ions around various groups of atoms provides detailed information of differences and similarities in their interactions with RNA molecules. With increasing concentration of Mg^2+^ ions, we find that K^+^ ions are readily displaced at specific binding sites near NBPOs. However even with high Mg^2+^ concentrations K^+^ ions retained certain binding sites around the RNA as well as contributing to the condensed layer of ions. Notably, ion condensation increases upon going from partially unfolded to the fully folded native state with the increase in ions in the condensed layer being supplied by K^+^ indicating that counter ion condensation by the monovalent ion contributes to stabilization of the tertiary structure. Localized K^+^-nucleobase interactions have a stabilizing effect on helical structure, including in partially unfolded states of Twister. Notably, in the presence of limited amounts of divalent ions, the monovalent ions may assume a supporting role in stabilizing the native, folded state of Twister by facilitating short NBPO pair interactions. Given Twisters ability to self-cleave at low Mg^2+^ concentrations further studies are required to address if these observations may be applied to other RNAs.

Many of the present results are consistent with those reported by Thirumalai and coworkers on alternate RNAs based on course-grained simulations in conjunction with both explicit ions (Denesyuk and Thirumalai 2015; Hori et al. 2019) and explicit divalent with a RISM monovalent ion model (Denesyuk et al. 2018; Nguyen et al. 2019). This includes the occurrence of RNA-Mg^2+^ interactions in unfolded states with the number of interactions increasing upon going to the folded state and increases in Mg^2+^ concentration increasing the extent of such interactions. The importance of specific Mg^2+^-RNA interactions is also observed. However, the present results indicate that as RNA folds the monovalent K^+^ ions also participate in charge neutralization, structural stability of secondary interactions, and sequence specific indications, even in the presence of high MgCl_2_, suggesting the importance of their explicit treatment in simulation studies versus the use of effective potentials. On the other hand, the small number of interactions of Mg^2+^ with the nucleobases indicates that the assumption in the course-grained model that omit direct Mg^2+^-base interactions is reasonable (Nguyen et al. 2019). Finally, the course-grained model is quite impressive in its ability to model the impact of ions on the thermodynamics of folding from fully unfolded states through the native states, a capability that is still a challenge for all-atom models. Thus, it is clear that both course-grained and all-atom based approaches are required to fully elucidate details on the folding of RNA, including the impact of ions.

## Materials and Methods

The Twister ribozyme, which exhibits a double pseudoknot comprised of strong secondary and tertiary interactions (Figure S2), was obtained from a crystallographic study (PDB ID: 4OJI) (Liu et al. 2014). The simulations were performed at four different concentrations of divalent ion (0, 10, 20 and 100 mM MgCl_2_) with a background buffer concentration of monovalent ions ranging from approximately 100 to 200 mM KCl with the number of ions and concentrations shown in Table S1 of the Supplementary Information. To sample both the native and partially unfolded states US was applied based on a RC defined as the distance between centers of masses of two groups of heavy atoms (Figure S1a). The native state corresponds to RC = 15 Å and the most unfolded conformation sampled corresponds to RC = 40 Å. In total, 26 windows were used along the RC for each of the four systems with each window being subjected to 2 sets of 5 cycles of GCMC/MD simulations. Each GCMC/MD cycle included six stages of GCMC sampling of ions and water molecules followed by 10 ns of production MD run. Overall, 2.6 microseconds of sampling for each of the four systems was performed using in-house code for GCMC (Lakkaraju et al. 2014; Sun et al. 2018) and OpenMM (version 7.1.1) for MD (Eastman et al. 2017). The first 4 ns of each 10 ns MD run were discarded as equilibration providing in total 1.56 microseconds of MD simulation per system to analyze the ion atmosphere around a range of RNA conformations. Snapshots from the trajectories were saved every 10 ps. Details of the simulation protocol and parameters can be obtained from the methods section of our previous study (Kognole and MacKerell 2020). Analysis of ion distributions included “grid free energies” (GFE) where GFE = −k_B_T ln(P), where P is the probability of the ion occupying a voxel (1 Å cubic unit of volume) on the RNA surface relative to the voxel occupancy of the same species in bulk solution, k_B_ is the Boltzmann constant, and T is the temperature (303.15 K)(Raman et al. 2013; Lemkul et al. 2016).

Analysis of ion count around Twister to characterize the ion atmosphere uses a cutoff scheme that was adopted in the present study to simplify the comparison between Mg^2+^ and K^+^ ions. Previously, in atomistic simulations a general cutoff of 9 Å associated with the Manning Radius based on Counter-Ion Condensation (CIC) theory has been used to separate the condensed ions from bulk ions when only monovalent ions are present in the system (Savelyev and MacKerell 2014). Recently, a coarse-grained approach calculated ions comprising the ion-atmosphere by defining a preferential interaction coefficient (Nguyen et al. 2019). Here, based on the radial distribution function (RDF) of Mg^2+^ and K^+^ ions around the NBPOs (Figures 7 and 8 below), we have implemented the following cutoffs to differentiate between direct inner-shell (0-3 Å), outer-shell (3-5.5 Å) and non-dehydrated (5.5-8 Å) interactions. Ions beyond 8 Å are considered outside of the condensed layer. The RDF of ions around a set of atoms was calculated by collecting the distances between the ions and the atoms over selected frames and calculating the probability of finding them at distance r. The probabilities are corrected for the change in volume of each shell used for counting the ions and further normalized with the number of selected RNA atoms and total number of frames. The ion counting analysis was performed by finding the total number of ions in spherical regions of given cutoff radius around the selected atoms. The counting was carried out over 5 sets of 12-ns trajectories at each reaction coordinate and errors were calculated as standard error of mean.

## Acknowledgements

We thank the National Institutes of Health [GM131710] for financial support for this work and the Computer-Aided Drug Design Center at the University of Maryland Baltimore for computing time.

## References

Cunha RA, Bussi G. 2017. Unraveling Mg 2+ –RNA binding with atomistic molecular dynamics. RNA 23: 628–638.

Denesyuk NA, Hori N, Thirumalai D. 2018. Molecular Simulations of Ion Effects on the Thermodynamics of RNA Folding. The Journal of Physical Chemistry B 122: 11860–11867.

Denesyuk NA, Thirumalai D. 2013. Coarse-Grained Model for Predicting RNA Folding Thermodynamics. The Journal of Physical Chemistry B 117: 4901–4911.

Denesyuk NA, Thirumalai D. 2015. How do metal ions direct ribozyme folding? Nat Chem 7: 793–801.

Disney MD. 2019. Targeting RNA with Small Molecules To Capture Opportunities at the Intersection of Chemistry, Biology, and Medicine. J Am Chem Soc 141: 6776–6790.

Draper DE. 2004. A guide to ions and RNA structure. RNA 10: 335–343.

Draper DE. 2008. RNA folding: thermodynamic and molecular descriptions of the roles of ions. Biophys J 95: 5489–5495.

Draper DE, Grilley D, Soto AM. 2005. Ions and RNA Folding. Annu Rev Bioph Biom 34: 221–243.

Eastman P, Swails J, Chodera JD, McGibbon RT, Zhao Y, Beauchamp KA, Wang LP, Simmonett AC, Harrigan MP, Stern CD et al. 2017. OpenMM 7: Rapid development of high performance algorithms for molecular dynamics. PLoS computational biology 13: e1005659.

Fischer NM, Poî Eto MD, Steuer J, Van Der Spoel D. 2018. Influence of Na + and Mg 2+ ions on RNA structures studied with molecular dynamics simulations. Nucleic Acids Research 46: 4872–4882.

Heilman-Miller SL, Pan J, Thirumalai D, Woodson SA. 2001a. Role of counterion condensation in folding of the Tetrahymena ribozyme. II. Counterion-dependence of folding kinetics. J Mol Biol 309: 57–68.

Heilman-Miller SL, Thirumalai D, Woodson SA. 2001b. Role of counterion condensation in folding of the Tetrahymena ribozyme. I. Equilibrium stabilization by cations. J Mol Biol 306: 1157–1166.

Hori N, Denesyuk NA, Thirumalai D. 2019. Ion Condensation onto Ribozyme Is Site Specific and Fold Dependent. Biophys J 116: 2400–2410.

Hu X, Provasi D, Ramsey S, Filizola M. 2019. Mechanism of μ-Opioid Receptor-Magnesium Interaction and Positive Allosteric Modulation DOI:10.1016/j.bpj.2019.10.007. Biophys J: In press.

Kim HD, Nienhaus GU, Ha T, Orr JW, Williamson JR, Chu S. 2002. Mg2+-dependent conformational change of RNA studied by fluorescence correlation and FRET on immobilized single molecules. Proc Natl Acad Sci U S A 99: 4284–4289.

Kirmizialtin S, Pabit Suzette A, Meisburger Steve P, Pollack L, Elber R. 2012. RNA and Its Ionic Cloud: Solution Scattering Experiments and Atomically Detailed Simulations. Biophys J 102: 819–828.

Kognole AA, MacKerell AD, Jr. 2020. Mg(2+) Impacts the Twister Ribozyme through Push-Pull Stabilization of Nonsequential Phosphate Pairs. Biophys J 118: 1424–1437.

Korman A, Sun H, Hua B, Yang H, Capilato JN, Paul R, Panja S, Ha T, Greenberg MM, Woodson SA. 2020. Light-controlled twister ribozyme with single-molecule detection resolves RNA function in time and space. Proceedings of the National Academy of Sciences: 202003425.

Lakkaraju SK, Raman EP, Yu W, Mackerell AD, Jr. 2014. Sampling of Organic Solutes in Aqueous and Heterogeneous Environments Using Oscillating Excess Chemical Potentials in Grand Canonical-like Monte Carlo-Molecular Dynamics Simulations. J Chem Theory Comput 10: 2281–2290.

Lemkul JA, Lakkaraju SK, Mackerell AD, Jr. 2016. Characterization of Mg 2+ Distributions around RNA in Solution. ACS Omega 1: 680–688.

Leonarski F, D’Ascenzo L, Auffinger P. 2019. Nucleobase carbonyl groups are poor Mg(2+) inner-sphere binders but excellent monovalent ion binders-a critical PDB survey. RNA 25: 173–192.

Leonarski F, D’Ascenzo L, Auffinger P. 2016. Binding of metals to purine N7 nitrogen atoms and implications for nucleic acids: A CSD survey. Inorganica Chimica Acta 452: 82–89.

Li W, Nordenskiöld L, Mu Y. 2011. Sequence-Specific Mg <sup>2+</sup> –DNA Interactions: A Molecular Dynamics Simulation Study. The Journal of Physical Chemistry B 115: 14713–14720.

Lippert B. 2000. Multiplicity of metal ion binding patterns to nucleobases. Coordination Chemistry Reviews 200-202: 487–516.

Liu Y, Wilson TJ, McPhee SA, Lilley DMJ. 2014. Crystal structure and mechanistic investigation of the twister ribozyme. Nat Chem Biol 10: 739–744.

MacKerell AD. 2019. Ions Everywhere? Mg2+ in the μ-Opioid GPCR and Atomic Details of Their Impact on Function. Biophysical Journal.

Manning GS. 1978. The molecular theory of polyelectrolyte solutions with applications to the electrostatic properties of polynucleotides. Quarterly Reviews of Biophysics 11: 179–246.

Nguyen HT, Hori N, Thirumalai D. 2019. Theory and simulations for RNA folding in mixtures of monovalent and divalent cations. Proceedings of the National Academy of Sciences 116: 21022.

Onoa B, Tinoco I, Jr. 2004. RNA folding and unfolding. Curr Opin Struc Biol 14: 374–379.

Pabit SA, Meisburger SP, Li L, Blose JM, Jones CD, Pollack L. 2010. Counting ions around DNA with anomalous small-angle X-ray scattering. J Am Chem Soc 132: 16334–16336.

Panja S, Hua B, Zegarra D, Ha T, Woodson SA. 2017. Metals induce transient folding and activation of the twister ribozyme. Nat Chem Biol 13: 1109–1114.

Peschke M, Blades AT, Kebarle P. 1998. Hydration Energies and Entropies for Mg2+, Ca2+, Sr2+, and Ba2+ from Gas-Phase Ion-Water Molecule Equilibria Determinations. The Journal of Physical Chemistry A 102: 9978–9985.

Raman EP, Yu W, Lakkaraju SK, MacKerell AD, Jr. 2013. Inclusion of multiple fragment types in the site identification by ligand competitive saturation (SILCS) approach. J Chem Inf Model 53: 3384–3398.

Savelyev A, MacKerell AD. 2014. Balancing the Interactions of Ions, Water, and DNA in the Drude Polarizable Force Field. The Journal of Physical Chemistry B 118: 6742–6757.

Shiman R, Draper DE. 2000. Stabilization of RNA tertiary structure by monovalent cations. J Mol Biol 302: 79–91.

Šponer J, Bussi G, Krepl M, Banáš P, Bottaro S, Cunha RA, Gil-Ley A, Pinamonti G, Poblete S, Jurečka P et al. 2018. RNA Structural Dynamics As Captured by Molecular Simulations: A Comprehensive Overview. Chemical Reviews 118: 4177–4338.

Šponer J, Sabat M, Gorb L, Leszczynski J, Lippert B, Hobza P. 2000. The Effect of Metal Binding to the N7 Site of Purine Nucleotides on Their Structure, Energy, and Involvement in Base Pairing. The Journal of Physical Chemistry B 104: 7535–7544.

Sun D, Lakkaraju K, Jo S, Mackerell AD, Jr. 2018. Determination of Ionic Hydration Free Energies with Grand Canonical Monte Carlo/Molecular Dynamics Simulations in Explicit Water. J Chem Theory Comput.

Thirumalai D, Lee N, Woodson SA, Klimov DK. 2001. Early Events in RNA Folding. Annual review of physical chemistry 52: 751–762.

Ucisik MN, Bevilacqua PC, Hammes-Schiffer S. 2016. Molecular Dynamics Study of Twister Ribozyme: Role of Mg <sup>2+</sup> Ions and the Hydrogen-Bonding Network in the Active Site. Biochemistry-US 55: 3834–3846.

Westhof E, Auffinger P. 2000. RNA Tertiary Structure. Encyclopedia of Analytical Chemistry: 5222–5232.

Woodson SA. 2010. Compact intermediates in RNA folding. Annu Rev Biophys 39: 61–77.

Xi K, Wang F-h, Xiong G, Zhang Z-l, Tan Z-j. 2018. Competitive Binding of Mg 2+ and Na + Ions to Nucleic Acids: From Helices to Tertiary Structures. Biophysical Journal 114: 1776–1790.

Yoo J, Aksimentiev A. 2012. Competitive Binding of Cations to Duplex DNA Revealed through Molecular Dynamics Simulations. The Journal of Physical Chemistry B 116: 12946–12954.

